# A negative role for transcription factor Bcr1 in *Candida albicans* hypha-associated gene expression

**DOI:** 10.64898/2026.09.03.749130

**Authors:** Anupam Sharma, Aaron P. Mitchell

**Author notes:** Correspondence: Aaron P. Mitchell. Anupam Sharma, Department of Molecular Microbiology and Immunology, Brown University, Providence, RI, 02906 USA.

## Abstract

Filamentation, a central virulence trait of the fungal pathogen *Candida albicans,* is required for host cell damage, tissue invasion, and biofilm formation. The transcription factor Bcr1 was first found to be required for biofilm formation but not filamentation. Subsequent studies revealed a negative role for Bcr1 in filamentation in opaque cells, a cell type that is required for mating, under hypoxic conditions, and in a *wor2*Δ/Δ mutant background. Here we characterize a new context in which Bcr1 negatively regulates filamentation and present its as sociated gene expression impact. We compared the wild-type and *bcr1*Δ/Δ mutant cells in the SC5314 reference strain background at 30°C in glucose media, conditions that do not induce filamentation. The *bcr1*Δ/Δ mutant displayed increased invasive growth in a solid medium, and some filamentation ability in a liquid medium. The *bcr1*Δ/Δ effect on filamentation was augmented by overexpression of hyphal cyclin gene *HGC1,* which caused pseudohyphal growth in wild-type, *efg1*Δ/Δ, or *brg1*Δ/Δ strains, yet caused hyphal growth in the *bcr1*Δ/Δ strain. RNA-sequencing (RNA-seq) analysis of 30°C cells shows that the *bcr1*Δ/Δ mutant has elevated expression of two genes that drive filamentation: *HGC1* and *UME6.* In published data these same two genes have decreased expression in the *bcr1*Δ/Δ mutant at 37°C, the temperature at which biofilm formation is typically assayed. A growing cadre of biofilm/hyphal regulators can have both positive and negative effects on hypha-associated gene expression, including Bcr1, Efg1, Ndt80, and Nrg1.

**IMPORTANCE:** Production of filamentous cells is a central virulence trait of the fungal pathogen *Candida albicans*. Evidence here shows that the transcription factor Bcr1 is a negative regulator of filamentation at low temperature (30°C). Bcr1 affects expression of two drivers of filamentation, the genes *HGC1* and *UME6.* Surprisingly, Bcr1 inhibits their expression at 30°C but stimulates their expression at 37°C. A growing number of filamentation regulators seem to function as both positive and negative regulators of filamentation, depending upon genetic and environmental contexts.

## INTRODUCTION

The fungal pathogen *Candida albicans* can grow as either yeast cells or filamentous cells, including hyphae and pseudohyphae, depending upon environmental conditions (1). Filamentation ability is required for virulence in an array of animal models (2, 3), and for biofilm formation (4), a source of infecting cells in human patients. Filamentation is also closely tied to the impact of antifungals (5). These connections have prompted significant interest in *C. albicans* filamentation and its control (6–8).

Given that filamentation is required for biofilm formation (4), it makes sense that many regulators of biofilm formation also govern filamentation. This connection arises from the transcriptional output that accompanies filamentation. Expression of a set of hypha-associated genes correlates with filamentous growth under many conditions (9–11). The precise members of this gene set vary with the conditions used to induce filamentation and those used to grow “control” yeast cells (9–11). However, one recent distillation of such several gene sets generated through RNA-seq yielded only 16 common hypha-associated genes (12), several of which contribute to biofilm formation (4). Those common genes include *HGC1* and *UME6*, for which increased expression is sufficient to drive filamentation and biofilm formation (13–17).

The transcription factor Bcr1 seemed like an exception to rule that biofilm regulators are filamentation regulators. Initial characterization indicated that a *bcr1*Δ/Δ mutant was defective in biofilm formation but not filamentation (18). This separation of phenotypes may have been a consequence of the range of growth conditions or strain backgrounds used (18, 19), because *bcr1*Δ/Δ mutants in other growth conditions and strain backgrounds are defective in filamentation (20). In addition, *bcr1*Δ/Δ mutants are defective in expression of several hypha-associated genes (18–23). Therefore, Bcr1 may be considered a conditional positive regulator of filamentation.

Several studies indicate that Bcr1 also functions as a negative regulator of filamentation. Guan et al. examined filamentation ability of opaque cells (24). Opaque cells have a critical role in *C. albicans* mating, and low temperature supports their stability and filamentation ability (25). Opaque cells that carry a *bcr1*Δ/Δ mutation showed elevated levels of filamentation in liquid or solid media at the low temperature of 25°C (24). Another study, by Desai et al., showed that a *bcr1*Δ/Δ mutant displayed increased filamentation under hypoxic conditions on solid medium at 25°C (26). In that context, Henry et al. have integrated Bcr1 into the broader network of hypoxia-responsive regulators (27). A third study, by Goerlich and Mitchell (23), looked at the genetic interaction between Bcr1 and the white-opaque regulator Wor2. At 37°C, double *bcr1*Δ/Δ *wor2*Δ/Δ mutants presented increased filamentation levels and increased expression of many hypha-associated genes, the same genes that have reduced expression in *bcr1*Δ/Δ single mutants. All of these studies indicate that Bcr1 has the capacity to negatively regulate filamentation, a capacity that depends upon specific environmental and genetic signals.

Our study here began with observations that indicate that Bcr1 negatively regulates filamentation in the SC5314 reference strain background under fairly conventional growth conditions at the low temperature of 30°C. RNA-seq analysis indicates that the hyphal drivers *HGC1* and *UME6* are repressed by Bcr1 at 30°C. The surprising twist is that these same two genes are activated by Bcr1 at 37°C.

## MATERIAL AND METHODS

### Strains and Media

All strains used in the work are listed in Supplemental Table S1. They were maintained in 15% glycerol stocks stored at -80°C. Strains were grown before all experiments on YPD medium (2% Bacto Peptone, 2% dextrose, 1% yeast extract) for 48 hrs at 30°C, and then cultured overnight in liquid YPD at 30°C with shaking. Transformants were selected on YPD + 400 μg/mL nourseothricin or SC-Arg (2% dextrose, 0.67% Difco yeast nitrogen base with ammonium sulfate without amino acids, 0.079% CSM-Arg mix, solidified with 2% agar). Filamentation assays were performed in YPD or liquid RPMI 1640 medium (Sigma-Aldrich), adjusted to pH 7.4 and supplemented with 10% fetal bovine serum (Atlanta Biologicals). Invasion assays were conducted on SC medium (2% dextrose,0.67% Difco yeast nitrogen base without amino acids, 0.079% CSM mix, solidified with 2% agar).

The strains constitutively expressing *HGC1* were constructed using a *NAT1-P_TDH3_* cassette. The cassette containing a flanking homology to the *HGC1* upstream region was amplified using primers “HGC1 OE F” and “HGC1 OE R” from plasmid pCJN542. All plasmids and primers used in the study are listed in Tables S2 and S3, respectively. SC5314 WT and mutant strains were transformed with 3 μg of this *NAT1-P_TDH3_* cassette, 1 μg of Cas9, and 1 μg of HGC1 P-2 sgRNA DNA cassette using a transient CRISPR-Cas9 system (28). The HGC1 P-2 sgRNA cassette was generated using split-joint PCR with primers “sgRNA/F HGC1P-2” and “SNR52/R HGC1P-2”. Transformants were selected on YPD + nourseothricin plates for the resistant phenotype. The genotyping was done by PCR using primers “HGC1 CH F” and “HGC1 CH R” for the presence of the native *HGC1* promoter, and “HGC1 CH F2” and “NAT CH R” for the presence of the *NAT1-P_TDH3_* cassette in the *HGC1* promoter region.

The *hgc1*Δ/Δ and *bcr1*Δ/Δ *hgc1*Δ/Δ double mutants were generated using wild-type and *bcr1*Δ/Δ strains obtained from the *C. albicans* transcription factor deletion library (29). The *HGC1* deletion cassette containing the *ARG4* selectable marker was amplified from plasmid pSN69 using primers HGC1 ARG4 DEL F and HGC1 ARG4 DEL R. The cassette was introduced into the parental strains using transient CRISPR-Cas9-mediated transformation (28), followed by selection on arginine-dropout medium. Homozygous *HGC1* deletion mutants were subsequently identified by PCR genotyping. An *ARG4* cassette was integrated into the genomes of wild-type and *bcr1*Δ/Δ strains to restore arginine prototrophy for phenotypic assays.

### Filamentation assays and imaging

The strains were inoculated to an OD_600_ of 0.5 from overnight cultures into 5 mL of YPD medium or RPMI supplemented with serum in glass culture test tubes. Cells were grown for 4 hours at 30°C or 37°C (as indicated in the text) in a roller drum for vigorous agitation. Cells were collected by centrifugation at 2800 rpm for 5 min and then fixed with 4% formaldehyde in PBS for 10 minutes with vortexing. Fixed cells were then washed twice in PBS and stained with Calcofluor-white (200 µg/mL in PBS). The imaging of stained cells was done using a slit-scan confocal optical unit on a Zeiss Axiovert 200 microscope.

### Agar invasion assay

The cells from the overnight culture were diluted to an OD_600_ of 1 in PBS. 3 μL of cells were spotted on the surface of a 10 cm SC agar plate at least 2 cm apart to prevent inhibition of growth due to nutrient depletion. Plates were incubated at 30°C for 6 to 8 days. The cells were removed from the surface of the plates by washing them with PBS. Images were acquired before and after the washing. For imaging of invading filaments, a block 1 mm wide was cut through the colony and agar with a razor blade. Then it was laid in a 12-well plate with 0.5 mL 100% Thiodiethanol. The cross-sectional images were taken with a 10X objective using a Keyence microscope.

### RNA extraction and sequencing

RNA extractions were done according to the previously described method (30). Briefly, strains were grown to log phase at 30°C and RNA extraction was performed using a Qiagen RNeasy mini kit (Cat#74104) with some modifications. RNA sequencing was performed as described in (31). Briefly, RNA-seq count data from three biological replicates each of wild-type (WT; SC5314) and *bcr1*Δ/Δ mutant cells grown in YPD medium at 30°C for 4 h were used for quality-control analysis. Gene-level count data were obtained from Dataset S1 and analyzed using DESeq2 (32). Genes with fewer than 10 total counts across all six samples were excluded, leaving 5,938 genes for analysis. Count normalization was performed using DESeq2 size-factor estimation, followed by variance-stabilizing transformation (VST) with blind = TRUE and a parametric dispersion fit. Pairwise Pearson correlation coefficients were calculated across all 5,938 retained genes using VST expression values to assess sample-to-sample similarity and reproducibility between biological replicates. No batch correction was applied.

### Software

Microscope Images were arranged, and adjustments were performed using Image J. All statistical analyses were carried out using GraphPad Prism, version 8.4.2. Gene Ontology (GO) term enrichment was performed using the Termfinder tool at the Candida Genome Database (33). Venn diagrams showing overlaps between gene sets were generated using Venny 2.1. RNA-seq quality-control analyses were initially performed using R software (version 4.3.3) and DESeq2 (version 1.42.0), and the variance-stabilizing transformation (VST) results were subsequently independently validated using R software (version 4.6.1) and DESeq2 (version 1.52.0). Volcano and heatmap plots were generated using python 3.12.1.

### Data Interpretation

Interpretations and hypotheses were informed by the extensive records available at the

*Candida* Genome Database (33).

### Data availability

All data necessary to support the conclusions of this study are available in this manuscript and our NCBI-deposited RNA-seq data sets with accession number GSE232192.

## RESULTS AND DISCUSSION

### Impact of Bcr1 on filamentation at 30°C

Two phenotypic assays indicated that Bcr1 may be a negative regulator of filamentation at 30°C in the *MTL***a**/*MTL*α reference strain SC5314. One assay was for agar invasion. In this assay cells were grown on SC medium at 30°C for 6 days. Agar invasion was assessed by retention of cells after washing the plate (Figure 1A), and by visualization of filament invasion in slices of the plate perpendicular to the plate surface (Figure 1A, side view). These growth conditions were noninducing or weakly inducing, as reflected by the modest agar invasion of the wild-type strain (Figure 1A, B). Control *efg1*Δ/Δ and *brg1*Δ/Δ mutations, which inactivate two biofilm/hyphal regulators, blocked invasion as expected. However, a *bcr1*Δ/Δ mutation increased invasion, as indicated by more extensive cell retention (Figure 1A) and deeper agar invasion (Figure 1A, side view) than the wild-type strain. The hyper-invasive *bcr1*Δ/Δ phenotype was reversed by complementation (Figure 1B), indicating that the *bcr1*Δ/Δ mutation causes the phenotype.

**Figure 1:**
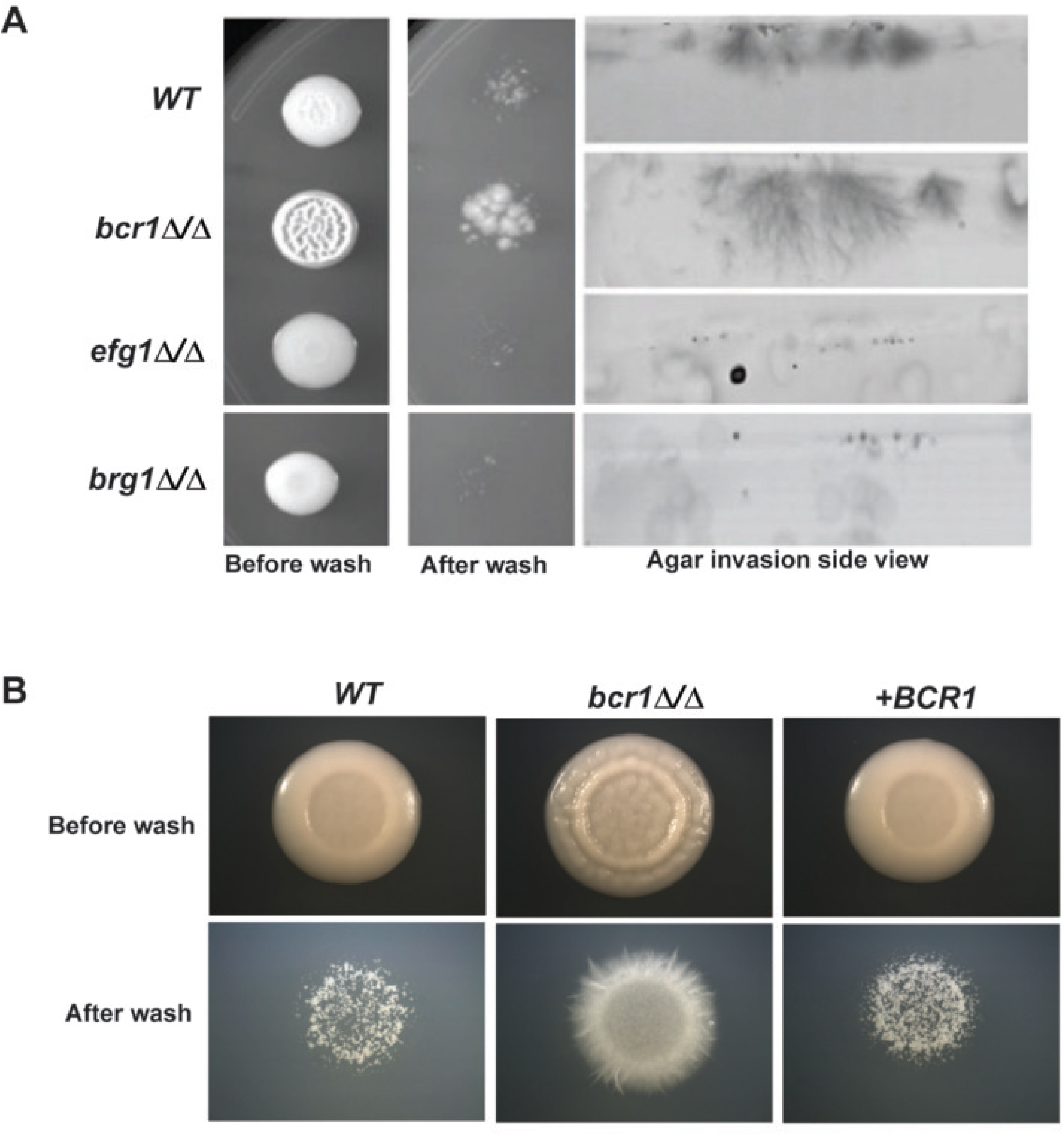
Agar invasion assays. **(A)** Representative agar-invasion assays done on SC agar at 30°C for 6 days comparing wild-type (WT), *bcr1*Δ/Δ, *efg1*Δ/Δ, and *brg1*Δ/Δ strains. Colonies were photographed before and after removal of surface-associated cells by washing. Side-view images show filament penetration into the agar. **(B)** WT, *bcr1*Δ/Δ, and the *BCR1*-complemented strain (+*BCR1*) were grown on SC agar at 30°C for 6 days and imaged before and after washing to remove non-invasive surface cells. Images are representative of independent experiments.

A second assay with convergent implications was of cell morphology. We used YPD medium at 30°C, a non-inducing condition. The wild-type strain produced mainly yeast cells (Figure 2A), as indicated by a length/width ratio of ∼1 (Figure 2B). The two control strains with *efg1*Δ/Δ or *brg1*Δ/Δ mutations also produced yeast cells. However, the strain that carried a *bcr1*Δ/Δ mutation produced a subset of elongated filamentous cells (Figure 2A, B). The effect was also evident qualitatively in derivatives of the strains that overexpress the hyphal cyclin gene *HGC1* (16). We fused both *HGC1* alleles to the *TDH3* promoter (*TDH3-HGC1*) in the four strains. In all cases, *TDH3*-*HGC1* caused abundant filamentation (Figure 2A, B), as expected (16). Filamentous cells in the wild-type, *efg1*Δ/Δ, and *brg1*Δ/Δ backgrounds were pseudohyphae because they lacked parallel sides and had constrictions at the septa between cells (Figure 2A). However, the filamentous cells in the *bcr1*Δ/Δ background resembled true hyphae because they had parallel sides with only minor constrictions at septa (Figure 2A). The *bcr1*Δ/Δ *TDH3-HGC1* cell units also had a significantly greater length/width ratio than the other *TDH3-HGC1* strains (Figure 2B). The stronger filamentation observed in the *bcr1*Δ/Δ *TDH3-HGC1* strain may reflect the combined effect of increased dosage of a downstream hyphal morphogenesis factor and loss of Bcr1-dependent repression at 30°C. These observations distinguish Bcr1 from Efg1 and Brg1, and indicate that Bcr1 functions as a negative regulator of filamentation under these assay conditions.

**Figure 2:**
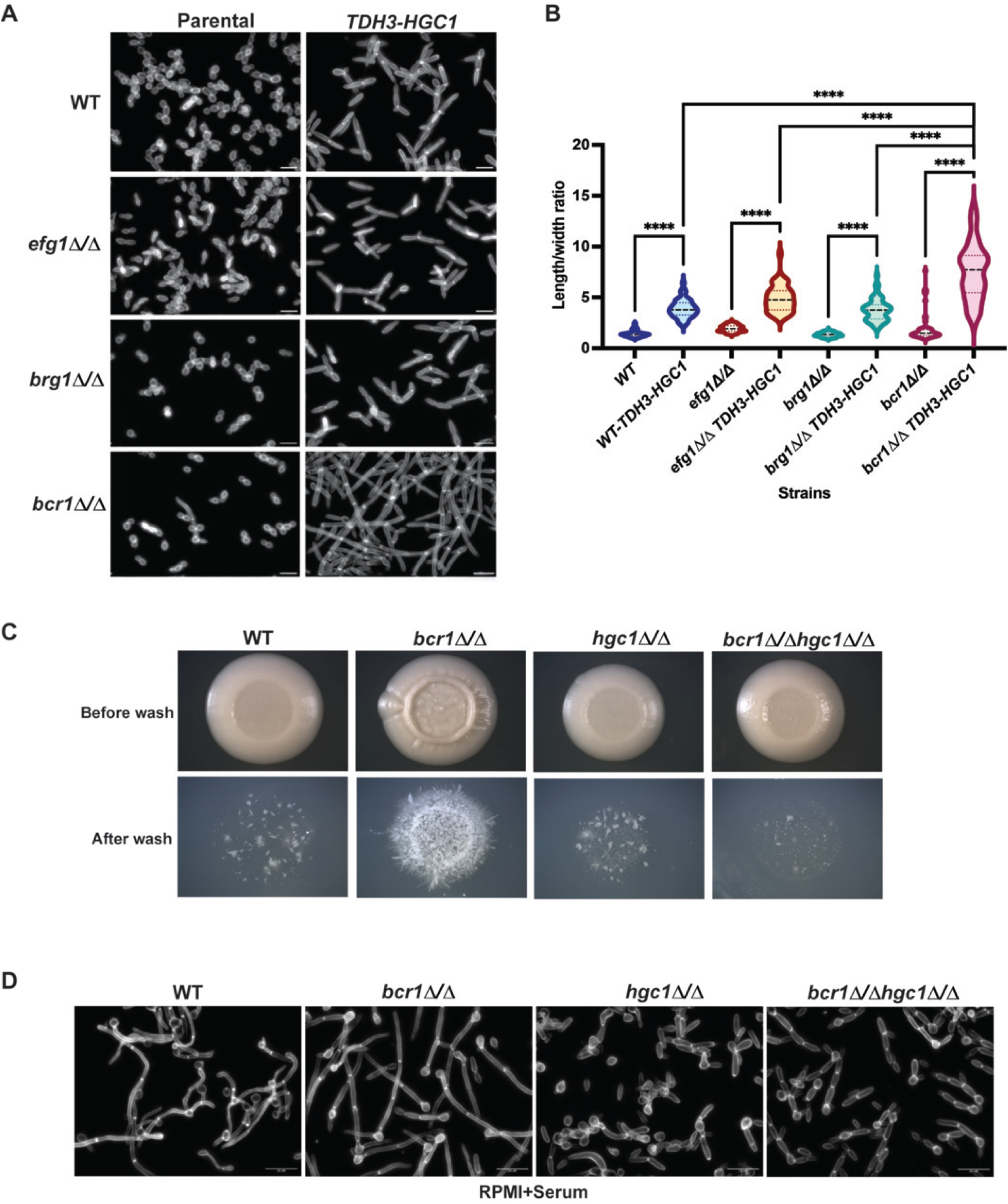
Impact of *HGC1* overexpression on filamentation. **(A)** Representative fluorescence images of parental and *TDH3-HGC1* strains in WT, *efg1*Δ/Δ, *brg1*Δ/Δ, and *bcr1*Δ/Δ genetic backgrounds. Constitutive *HGC1* expression promoted cellular elongation in each genetic background, with the strongest filamentation observed in *bcr1*Δ/Δ *TDH3-HGC1* cells. Scale bars, 5 µm**. (B)** Violin plots showing cellular length-to-width ratios for the indicated strains; 100–150 cells were analyzed per strain. Internal lines indicate the median and quartiles. Statistical comparisons were performed using Sidak’s multiple-comparisons test and are indicated by brackets; ****, *P* < 0.0001. **(C)** Representative agar invasion assays of wild-type (WT), *bcr1*Δ/Δ, *hgc1*Δ/Δ, and *bcr1*Δ/Δ *hgc1*Δ/Δ strains grown on SC agar at 30°C for 8 days. Colonies were photographed before (top) and after (bottom) washing to remove noninvasive surface cells. Images are representative of independent experiments. **(D)** Representative fluorescence images of WT, *bcr1*Δ/Δ, *hgc1*Δ/Δ, and *bcr1*Δ/Δ *hgc1*Δ/Δ strains grown in serum-supplemented RPMI medium for 4 h at 37°C. Scale bars, 20 µm.

To assess the role of Hgc1 in the filamentation phenotype of the *bcr1*Δ/Δ mutant, we examined a *bcr1*Δ/Δ *hgc1*Δ/Δ double mutant. Previous studies have established that Hgc1 is required for hyphal elongation, with *hgc1*Δ/Δ mutants exhibiting defective hyphal extension and shortened filaments (14, 17). On SC agar at 30°C, both the *hgc1*Δ/Δ single mutant and the *bcr1*Δ/Δ *hgc1*Δ/Δ double mutant exhibited defective agar invasion compared with the *bcr1*Δ/Δ mutant (Figure 2C). Similarly, following growth in RPMI medium supplemented with serum at 37°C for 4 h, the *hgc1*Δ/Δ mutant displayed impaired hyphal elongation, and the double mutant exhibited similarly reduced filamentation compared with the *bcr1*Δ/Δ mutant (Figure 2D). These findings indicate that Hgc1 is required for the enhanced filamentation and invasive growth associated with loss of Bcr1.

### Bcr1-responsive gene expression at 30°C

Bcr1-responsive gene expression at 37°C has been extensively characterized (18–23, 34). However, we are unaware of any genome-wide analysis of Bcr1-responsive gene expression with cells grown at 30°C, conditions that might reveal the gene regulatory basis for negative control of filamentation by Bcr1. We conducted RNA-seq analysis of SC5314 wild-type and *bcr1*Δ/Δ cells grown at 30°C in YPD medium for 4 h. Quality-control analysis of the 30°C RNA-seq dataset supported by pairwise Pearson correlations, including WT replicate correlations of 0.983-0.993 and *bcr1*Δ/Δ replicate correlations of 0.992-0.994 (Figure S1A-B, Dataset S1). We defined Bcr1-responsive genes with conventional cutoffs of a P_adj_ < 0.05 and a |log_2_ fold change| >1 in for the *bcr1*Δ/Δ mutant vs wild type comparison. We found 120 genes that were differentially expressed in the *bcr1*Δ/Δ vs wild type comparison (Dataset S1, Figure 3A). The 75 upregulated genes and 45 downregulated genes were each enriched for Gene Ontology (GO) terms related to the cell wall and cell surface (Dataset S1), as expected from Bcr1-responsive genes previously described in cells grown at 37°C (18–23, 34).

**Figure 3:**
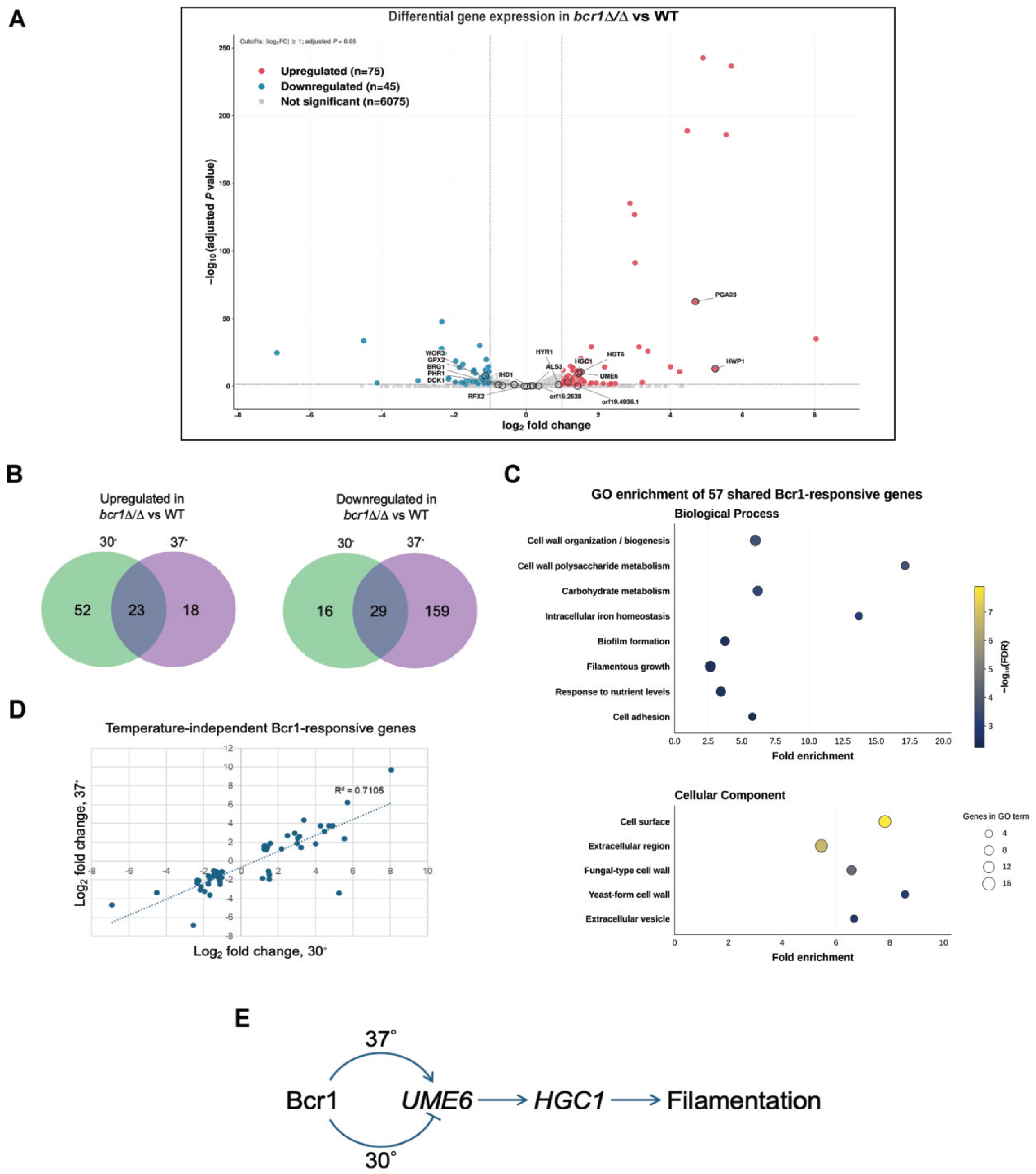
Bcr1 regulation of temperature-dependent and -independent transcriptional programs. **(A)** Volcano plot showing differential gene expression in *bcr1*Δ/Δ cells relative to WT cells grown at 30°C. RNA-seq analysis was performed using three biological replicates per strain. The x-axis shows log₂ fold change, and the y-axis shows −log₁₀-adjusted *P* value. Genes with an absolute log₂ fold change >1 and an adjusted *P* value <0.05 were classified as differentially expressed. Upregulated genes are shown in magenta, downregulated genes in cyan, and nonsignificant genes in gray. Sixteen genes associated with filamentation or Bcr1-dependent regulation are labeled. *BCR1* and *HIS1* were excluded from the downregulated gene count displayed in the plot, leaving 45 downregulated genes. **(B)** Venn diagrams comparing genes upregulated or downregulated in *bcr1*Δ/Δ relative to WT at 30°C with previously identified Bcr1-responsive genes at 37°C. **(C)** Bubble plots showing representative enriched Biological Process and Cellular Component Gene Ontology (GO) terms among the 57 genes differentially expressed in *bcr1*Δ/Δ relative to wild-type cells at both temperatures. Bubble size represents the number of genes associated with each GO term, color indicates −log₁₀ false discovery rate (FDR), and the x-axis shows fold enrichment. Only significantly enriched GO terms (FDR < 0.05) are shown. Complete enrichment results are provided in Dataset S1. **(D)** Comparison of log₂ fold changes at 30°C and 37°C for the 30°C/37°C Bcr1-responsive genes. A dotted regression line is included. **(E)** Proposed relationships that mediate temperature-dependent Bcr1 regulation. Bcr1 negatively regulates expression of *UME6* and *HGC1* at 30°C; Bcr1 positively regulates expression of *UME6* and *HGC1* at 37°C.

Our group recently published an RNA-seq analysis of a *bcr1*Δ/Δ mutant and wild-type strain grown at 37°C in YPD for 4 hours (23). Data in the current study were generated with the same strains grown at 30°C in YPD for 4 hours. The two datasets offer an opportunity to compare Bcr1-responsive genes at the two temperatures (Figure 3B). Of the 75 genes upregulated in the *bcr1*Δ/Δ mutant at 30°C, 23 genes were also upregulated at 37°C. Of the 45 genes downregulated in the *bcr1*Δ/Δ mutant at 30°C, 29 genes are also downregulated at 37°C. Overall, more genes are upregulated in the mutant at 30°C; fewer genes are downregulated in the mutant at 30°C (Figure 3B). Many transcription factors exert both positive and negative regulatory effects, and temperature may influence Bcr1 activity. *BCR1* expression was approximately 2.1-fold higher at 37°C than at 30°C (mean FPKM: 66.97 versus 31.84), potentially contributing to its temperature-dependent regulatory effects.

We defined a set of 30°C/37°C Bcr1-responsive genes, comprising the genes that are significantly regulated by Bcr1 at both temperatures (Dataset S1). The 57 genes in this set are enriched for cell wall-related GO terms in both the Biological Process and Cellular Component categories (Dataset S1, Figure 3C). The log_2_ fold change values at the two temperatures correlated with an R^2^ = 0.71 (Figure 3D). Only 5 genes had opposite fold changes at the two temperatures: *HWP1, HGT6, GCA1, HGC1*, and *UME6*. These genes are all upregulated in the *bcr1*Δ/Δ mutant at 30°C and downregulated in the *bcr1*Δ/Δ mutant at 37°C. Four of the five genes are within the hypha-associated gene set (*HWP1, HGT6, HGC1*, and *UME6*) distilled from multiple studies (12). (We note that only a portion of a broader hypha-associated gene set (10) was upregulated (Figure S2), an indication that the *bcr1*Δ/Δ transcriptional response was selective.) We examined prior genome-wide binding datasets to determine whether these five genes are likely direct Bcr1 targets. In the Bcr1 ChIP-chip dataset from Nobile et al. (19), only *UME6* was identified a Bcr1-bound target. In the Ume6 ChIP-seq dataset from Do et al. (35), *HWP1* and *HGC1* were identified as Ume6-bound targets. Thus, these findings suggest that Bcr1 may directly regulate *UME6*, while its effects on the other genes are likely indirect (Figure 3F). (We acknowledge explicitly the alternative possibility that Bcr1 may regulate *UME6* expression indirectly.) Increased expression of each of the genes *HGC1* and *UME6* can drive hypha formation under noninducing conditions, including low temperature (13–17). We propose that the reversal in regulatory impact of a *bcr1*Δ/Δ mutation on these two genes causes the *bcr1*Δ/Δ mutation to have opposite effects on filamentation at the two temperatures.

Many synopses of the regulation of filamentation and biofilm formation depict key transcription factors as either positive or negative regulators of hypha-associated genes and their biological consequences. This study emphasizes a more nuanced view that dates back to the discovery that Efg1, while a positive regulator of filamentation at 37°C (36), is a negative regulator of filamentation under embedded growth conditions at low temperature (37). Temperature and genetic background have been shown to reverse the gene expression impact of such well-known regulators as Efg1 (20, 38), Bcr1 (23), Brg1 (20), Ndt80 (38), and Nrg1 (39). Bcr1 is a negative regulator of filamentation under several conditions (this study and (23, 24, 26)). Gene expression impact of a *bcr1*Δ/Δ mutation has been assayed both low temperature (as reported here) or in a *wor2*Δ/Δ mutant background (23). Interestingly, the same five genes – *HWP1, HGT6, GCA1, HGC1*, and *UME6* – are negatively regulated by Bcr1 in both contexts, yet positively regulated by Bcr1 in an otherwise wild-type background at 37°C (this study and (23)). We speculate that the same mechanism may be triggered by the two contexts, low temperature or a *wor2*Δ/Δ defect, to explain the reversal of Bcr1 impact on these five genes. It is known that Bcr1 can form phase-separated complexes with other biofilm/hyphal regulators (40). Such complexes can vary in their precise composition, depending on expression levels of constituent proteins (41, 42). Perhaps low temperature or a *wor2*Δ/Δ defect restructure Bcr1 complexes to alter their regulatory impact. Alternatively, other protein complexes at the promoter regions of the five affected genes may be functionally altered in response to temperature changes.

The understanding that *C. albicans* Bcr1 has dual roles in hypha-associated gene expression offers some cohesion with observations in other *Candida* species. The Bcr1 ortholog in the emerging pathogen *Candida auris* is a negative regulator of filamentation (43). The negative impact of the Bcr1 ortholog, called Gfc1, is manifested at low temperature (25°C). A sampling of *C. auris* candidate genes showed that the Gfc1 negatively regulates *UME6* and *HGC1*. An exciting though speculative possibility is that the close connections among temperature, Bcr1, Ume6, and Hgc1 will be a common feature of filamentation control in *Candida* species.

## ACKNOWLEDGMENTS

We are grateful to Drs. Eunsoo Do, Katharina Goerlich, Yinhe Mao, Liping Xiong, Min-Ju Kim, Max Cravener, Amelia White, and Fred Lanni for their continued interest and suggestions. We thank Max Kuhr for outstanding laboratory management.

