## Supplemental table 3 for "A negative role for transcription factor Bcr1 in *Candida albicans* hypha-associated gene expression"

**Supplementary Table S3**

| Primer name | Sequence |
|---|---|
| **sgRNA/F HGC1** | GGTATCGATACCGATAATGAGTTTTAGAGCTAGAAATAGCAAGT TAAA |
| **SNR52/R HGC1** | TCATTATCGGTATCGATACCCAAATTAAAAATAGTTTACGCAAGTC |
| **SNR52/F** | AAGAAAGAAAGAAAACCAGGAGTGAA |
| **sgRNA/R** | ACAAATATTTAAACTCGGGACCTGG |
| **SNR52/N** | GCGGCCGCAAGTGATTAGACT |
| **sgRNA/N** | GCAGCTCAGTGATTAAGAGTAAAGATGG |
| **CaCas9/for** | ATCTCATTAGATTTGGAACTTGTGGGTT |
| **CaCas9/rev** | TTCGAGCGTCCCAAAACCTTCT |
| **pNAT F** | TTTCCCAGTCACGACGTT |
| **NAT1 CHR** | GTTCTGTATCTATAAGCAGTATCATCCAAAGTAGT |
| **HGC1 OE F** | CCCAAACTATACTTCCCAATAAAAGATAGAAACTCGCTTACAACAACACAATCC TGAAGATTATTAAATCTCTAATTTTCATCAAGCTTGCCTCGTCCCC |
| **HGC1 OE R** | ATGTTTTTGTATGGATGTTGTTGTTGTTGTTTTTGTTGTGAAATTGATTTTGGAG TTAATGGTTTAGTTATATTTATCATtgttaatTAATTTGATTGTAAAGTTTGTTGATG |
| **sgRNA/F HGC1P-2** | GTGTGTATAGTGTAGTATCCGTGTTTTAGAGCTAGAAATAGCAAGTTAAA |
| **SNR52/R HGC1P-2** | ACGGATACTACACTATACACACCAAATTAAAAATAGTTTACGCAAGTC |
| **HGC1 CH F** | CTTACATTTTAGACGACCAACGGATACTACA |
| **HGC1 CH R** | CTTCGATTGAAGGATCATTTAAAGACCATTCTAAA |
| **HGC1 CH F2** | ACTCTCTTGTTGTTGTTGTTGTTGTTTATC |
| **HGC1 ARG DEL F** | ATCAAAGCATCAAACCAATACCCAACACTTTAATATCTAGGGTTTCCATTCACA TATACACATATAAACATATATTAATAaagaacatttctgtaccgcaaatgt |
| **HGC1 ARG DEL R** | AAAACATCATTAAAATTTCATATCATAATAACAACATCTTTCTCCATTCTCCATTC TCTACTTTATCTTTCTCTCTTTCTacaaaagctatttgcatcgtttgaaag |

**Table S3: List of primers and their nucleotide sequence.**
