## Supplemental table 2 for "A negative role for transcription factor Bcr1 in *Candida albicans* hypha-associated gene expression"

### Supplementary Table S2

| Plasmid name | Description | Marker | Reference |
|---|---|---|---|
| pNAT | *NAT1* marker | $Amp^R$ | Min *et al.*, 2016 |
| pV1093 | CaCas9/sgRNA expression vector | $Amp^R$ | Vyas *et al.*, 2015 |
| pCJN542 | *NAT1-TDH3* promoter | $Amp^R$ | Nobile *et al.*, 2008 |
| pSN69 | *C. dubliniensis ARG4* gene deletion cassette | $Kan^R$ | Noble and Johnson, 2005 |

**Table S2: Plasmids used in this study**
