## Supplemental table 1 for "A negative role for transcription factor Bcr1 in *Candida albicans* hypha-associated gene expression"

### Supplementary Table S1

| Strain no. | Strain | Species | Genotype/Parent |
|---|---|---|---|
| ASM1 | SC5314 | *C.albicans* | Wild type clinical isolate |
| ASM329 | SC5314 *efg1Δ/Δ* | *C.albicans* | *efg1Δ::r1HIS1r1/efg1Δ::r1HIS1r1 his1Δ::r3/his1Δ::r3* |
| ASM331 | SC5314 *bcr1Δ/Δ* | *C.albicans* | *bcr1Δ::r1HIS1r1/bcr1Δ::r1HIS1r1 his1Δ::r3/his1Δ::r3* |
| ASM338 | SC5314 *brg1Δ/Δ* | *C.albicans* | *brg1Δ::r1HIS1r1/brg1Δ::r1HIS1r1 his1Δ::r3/his1Δ::r3* |
| ASM350 | SC5314 *efg1Δ/Δ TDH3-HGC1* | *C.albicans* | *efg1Δ::r1HIS1r1/efg1Δ::r1HIS1r1 HGC1:: NAT1-$P_{TDH3}$-HGC1/HGC1:: NAT1-$P_{TDH3}$-HGC1 his1Δ::r3/his1Δ::r3* |
| ASM355 | SC5314 *brg1Δ/Δ TDH3-HGC1* | *C.albicans* | *brg1Δ::r1HIS1r1/brg1Δ::r1HIS1r1 HGC1:: NAT1-$P_{TDH3}$-HGC1/HGC1:: NAT1-$P_{TDH3}$-HGC1 his1Δ::r3/his1Δ::r3* |
| ASM363 | SC5314 WT *TDH3-HGC1* | *C.albicans* | *HGC1::NAT1-$P_{TDH3}$-HGC1/ HGC1:: NAT1-$P_{TDH3}$-HGC1* |
| ASM372 | SC5314 *bcr1Δ/Δ TDH3-HGC1* | *C.albicans* | *bcr1Δ::r1HIS1r1/bcr11Δ::r1HIS1r1 HGC1:: NAT1-$P_{TDH3}$-HGC1/HGC1:: NAT1-$P_{TDH3}$-HGC1 his1Δ::r3/his1Δ::r3* |
| MH349 | *+BCR1* | *C.albicans* | *bcr1Δ::$BCR1^{SC5314}$-NAT1/bcr1Δ::r1HIS1r1 his1Δ::r3/his1Δ::r3* |
| ASM388 | WT | *C.albicans* | *leu2Δ/leu2Δ, ura3Δ/URA3, his1Δ/his1Δ, arg4Δ:ARG4/arg4Δ, iro1Δ/IRO1* |
| ASM712 | *bcr1Δ/Δ* | *C.albicans* | *bcr1Δ::C.d.HIS1/bcr1Δ::C.m.LEU2, leu2Δ/leu2Δ, ura3Δ/URA3, his1Δ/his1Δ, arg4Δ:ARG4/arg4Δ, iro1Δ/IRO1* |
| ASM700 | *hgc1Δ/Δ* | *C.albicans* | *hgc1Δ:ARG4/hgc1Δ:ARG4, leu2Δ/leu2Δ, ura3Δ/URA3, his1Δ/his1Δ, arg4Δ/arg4Δ, iro1Δ/IRO1* |
| ASM704 | *bcr1Δ/Δ hgc1Δ/Δ* | *C.albicans* | *hgc1Δ:ARG4/hgc1Δ:ARG4 bcr1Δ::C.d.HIS1/bcr1Δ::C.m.LEU2, leu2Δ/leu2Δ, ura3Δ/URA3, his1Δ/his1Δ, arg4Δ/arg4Δ, iro1Δ/IRO1* |

**Table S1: *Candida albicans* strains used in this study**
