## Supplemental figure 2 for "A negative role for transcription factor Bcr1 in *Candida albicans* hypha-associated gene expression"

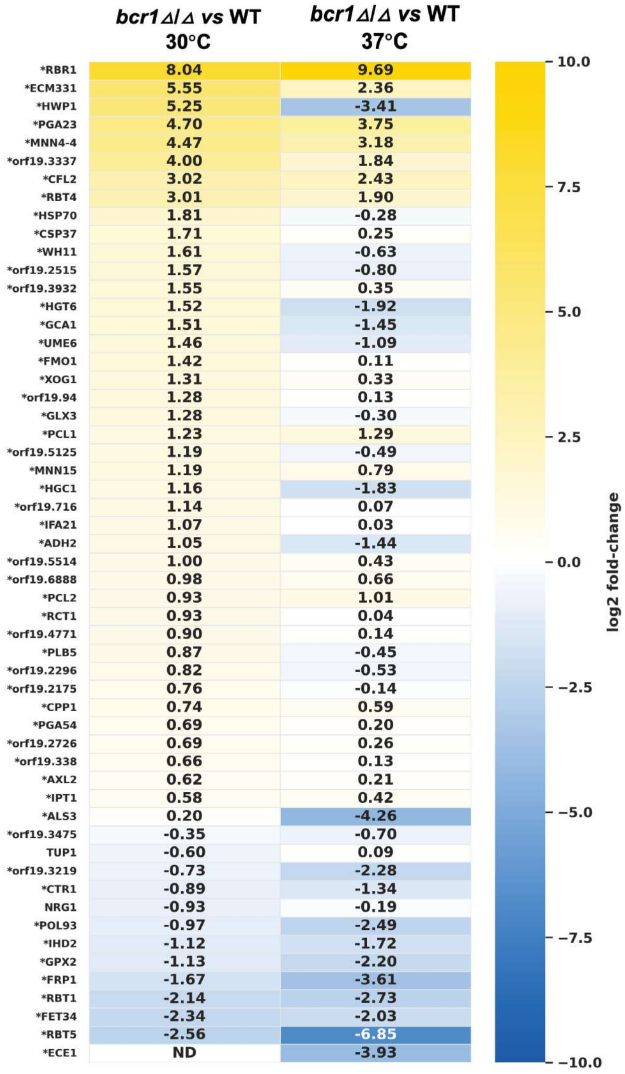


**Figure S2:** Heatmap showing $\log_2$ fold changes in hypha-associated genes defined by Cravener et al. (2023) in *bcr1*Δ/Δ relative to wild-type (WT) cells grown at 30°C and 37°C. Yellow indicates increased expression, blue indicates decreased expression, and white indicates little or no change. Values represent $\log_2$ fold changes, and asterisks indicate genes differentially expressed at 30°C (adjusted P < 0.05 and $|\log_2$ fold change$| \geq 1$). ND (expression not detected).
