## Supplemental figure 1 for "A negative role for transcription factor Bcr1 in *Candida albicans* hypha-associated gene expression"

**Figure S1:**

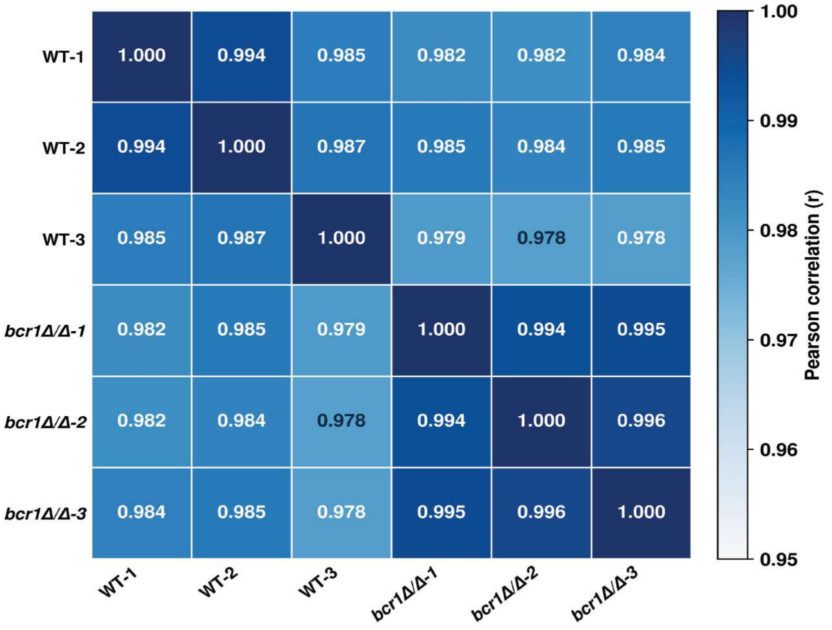


**Figure S1: Pearson correlations of RNA-seq data from wild-type and *bcr1*Δ/Δ cells.** RNA-seq analysis from wild-type (WT) and *bcr1*Δ/Δ mutant cells grown in YPD medium at 30°C for 4 h, with three independent biological replicates per genotype. Heatmap of pairwise Pearson correlation coefficients calculated from VST expression values of 5,938 genes across all six biological samples. Correlation coefficients are displayed within individual cells. All samples exhibited high pairwise correlations (r = 0.978–0.996 for distinct sample pairs), indicating overall similarity in genome-wide expression profiles.
